# *‘%svy_logistic_regression*: A generic SAS® macro for simple and multiple logistic regression and creating quality publication-ready tables using survey or non-survey data

**DOI:** 10.1101/575605

**Authors:** Jacques Muthusi, Samuel Mwalili, Peter Young

**Affiliations:** Division of Global HIV and Tuberculosis, U.S. Centers for Disease Control and Prevention, Nairobi, Kenya

**Keywords:** SAS macro, odds ratio, logistic regression, nutrition survey, data reporting, reproducible research

## Abstract

**Introduction:** Reproducible research is increasingly gaining interest in the research community. Automating the production of research manuscript tables from statistical software can help increase the reproducibility of findings. Logistic regression is used in studying disease prevalence and associated factors in epidemiological studies and can be easily performed using widely available software including SAS, SUDAAN, Stata or R. However, output from these software must be processed further to make it readily presentable. There exists a number of procedures developed to organize regression output, though many of them suffer limitations of flexibility, complexity, lack of validation checks for input parameters, as well as inability to incorporate survey design.

**Methods:** We developed a SAS macro, ***%svy_logistic_regression***, for fitting simple and multiple logistic regression models. The macro also creates quality publication-ready tables using survey or non-survey data which aims to increase transparency of data analyses. It further significantly reduces turn-around time for conducting analysis and preparing output tables while also addressing the limitations of existing procedures.

**Results:** We demonstrate the use of the macro in the analysis of the 2013-2014 National Health and Nutrition Examination Survey (NHANES), a complex survey designed to assess the health and nutritional status of adults and children in the United States. The output presented here is directly from the macro and is consistent with how regression results are often presented in the epidemiological and biomedical literature, with unadjusted and adjusted model results presented side by side.

**Conclusions:** The SAS code presented in this macro is comprehensive, easy to follow, manipulate and to extend to other areas of interest. It can also be incorporated quickly by the statistician for immediate use. It is an especially valuable tool for generating quality, easy to review tables which can be incorporated directly in a publication.

## Introduction

The principles of reproducible research are increasingly gaining interest both in the research community (1–5) and in the popular imagination as a result of high-profile failures to reproduce results. While funders and journals are increasingly requiring both publications and their supporting data be made publicly available with few exceptions (6–8) there has been less focus on the reproducibility of the analysis process itself. Reproducible research refers to increasing the transparency of the research endeavor by making the initial data, detailed analysis steps, and tools available to allow others to reproduce ones’ findings. Peng and Leek refer to increasing reproducibility as a tool to reduce the time required to uncover errors in analysis (9). One important link in the reproducible research value chain is eliminating manual reformatting of results from statistical software into draft manuscript tables.

In most epidemiological studies, one of the main outcomes of interest is disease prevalence – i.e. the proportion of all study subjects with a disease. Researchers are often interested in the probability or odds of subjects having a disease as well as associated predictive factors. These factors can be categorical (such as gender), ordinal (for example age categories), or continuous (for instance duration on treatment). The measures of association are often presented as crude (unadjusted) odds ratios from simple logistic regression or they can be presented as adjusted odds ratios from multiple logistic regression. In scientific reports from observational epidemiological studies it is common to combine the results from multiple statistical models and present the odds ratios side by side in complex tables showing the association between multiple covariates with the outcome of interest, both unadjusted and adjusted.

Logistic regression models can be fitted easily using available standard statistical analysis software such as SAS, SUDAAN, Stata or R, among others, and have been extended to handle weights and/or specialized variance estimation to account for complex survey designs. However, output from these software is not formatted for use directly in a publication and must be re-organized in order to make it more presentable based on the cultural norms of the biomedical literature or the specific requirements of the scientific journal (10, 11). Most epidemiological publications present regression tables showing odds ratios estimates and the corresponding 95% confidence intervals and/or p-values. They further enrich the output by including frequencies and proportion of study subjects who experienced the outcome of interest. Results from simple and multiple regression can also be presented side-by-side in one table. Some examples of publications which adopt this convention of presenting regression results are provided in the references (12–15). In order to accomplish this, one has to manually copy different parts of output into a template. This is both time-consuming and potentially prone to errors when revisions to the analysis are required.

A number of programs have been developed to facilitate conversion of regression output from statistical analysis software into formatted tables for publications. In Stata, several programs including ***esttab*** (16, 17), ***reformat*** (18), ***outreg2*** (19) are useful in formatting regression output. In R such packages as ***stargazer*** (20), ***broom*** (21), ***flextable*** (22) have also being found helpful. Though they are useful to statisticians they suffer from numerous limitations. For instance, they cannot automatically combine results from several simple logistic regression into a single table. It is also not possible to combine results from simple and multiple logistic regression into one output table. They are also not fully generic in that one has to explicitly specify variable labels and levels of categorical variables instead of extracting these from metadata. Further manipulation of output, for example, concatenating odds ratios and the corresponding 95% confidence interval into one column cell, has to be done manually which increases the risk of typographical errors in the output table. In SAS software, logistic regression models can be fitted using the LOGISTIC, GENMOD and SURVEYLOGISTIC procedures (23), though output from these procedures must be formatted further to make it presentable. SAS provides a flexible and powerful macro language that can be utilized to create and populate numerous table templates for presenting regression results. However, limited programming work has being done in SAS to date. There are several macros including ***%table1*** (24), ***%logistic_table*** (25) and ***%UniLogistic*** (26) which have been developed to assist in processing the output from regression procedures, but they are largely limited in terms of flexibility, lack of support for complex survey designs, or are unable to incorporate both categorical and continuous variables in one macro call. For instance, the macro, ***%table1***, presents variable names instead of the more meaningful variable labels. The other macros, ***%logistic_table***, and ***%UniLogistic***, produce output from simple logistic regression but not from multiple logistic regression. Also the ***%UniLogistic*** macro does not accommodate survey design parameters. Furthermore, these macros lack validation checks for input parameters and also do not export the output into word processing and spreadsheet programs for ease of incorporating into a publication.

## Methods

Recognizing the limitations of existing tools, we have designed a SAS macro, ***%svy_logistic_regression*** to help overcome these shortcomings while supporting the principles of reproducible research. The macro specifically organizes output from SAS procedures and formats it into quality publication epidemiologic tables containing regression results. We describe the macro functionality and provide an example analysis of a publicly-available dataset and provide access to the source code for the macro to allow others to use and extend it to support their own reproducible research.

Our developed SAS macro allows for both simple and multiple logistic regression analysis. Moreover, this SAS macro combines the results from simple and multiple logistic regression analysis into a single quality publication-ready table. The layout of the resulting table is consistent with how models are often presented in the epidemiological and biomedical literature, with unadjusted and adjusted model results presented side by side.

The macro, written in SAS software version 9.3 (27), runs logistic regression analysis in a sequential and interactive manner starting with simple logistic regression models followed by multiple logistic regression models using SAS PROC SURVEYLOGISTIC procedure. Frequencies and totals are obtained using PROC SURVEYMEANS and PROC SURVEYFREQ procedures. The final output is then processed using PROC TEMPLATE, PROC REPORT procedures and the output delivery system (ODS).

The macro is made up of six sub-macros. The first sub-macro, ***%svy_unilogit***, fine-tunes the dataset by applying the conditional statements, and computing the analysis domain size, thus preparing a final analysis dataset. It also prepares the class variables and associated reference categories. It calls the second and third sub-macros, ***%svy_logitc*** and ***%svy_logitn***, to perform separate simple (survey) logistic regression model on each categorical or continuous predictor variable respectively. It further processes results outputs into one table. The fourth sub-macro, ***%svy_multilogit***, performs multiple (survey) logistic regression on selected categorical and continuous predictor variables and processes result outputs into one table. The fifth sub-macro, ***%svy_printlogit***, combines results from ***%svy_unilogit*** and ***%svy_multilogit*** sub-macros and processes the output into an easy to review table which is exported into Microsoft word processing and excel spreadsheet programs. In addition, where survey design variables have been specified the macro automatically incorporates them into the computation. The sixth sub-macro, ***%runquit***, is executed after each SAS procedure or DATA step, to enforce in-build SAS validation checks on the input parameters. These include but not limited to checking if the specified dataset exists, ensuring required variables are specified, and verifying that values for reference categories for outcome, domain and categorical variables exist and are valid, as well as checking for logical errors. Once an error is encountered, the macro stops further execution and prints the error message on the log for the user to address it.

The macro is generic in that it can be used to analyze any dataset intended to fit a logistic regression model from survey or non-survey settings. It accepts both categorical and continuous predictor variables. Where survey data are used, it allows one to specify design-specific variables such as strata, clusters or weights. Domain analysis for sub-population estimation is also provided for by the macro. Ignoring domain analysis and instead performing a subset analysis will lead to incorrect standard errors. For non-survey settings, the survey input parameters like weights and cluster are set to a default value of 1.

The macro also allows the user to explicitly specify the level or category of the binary outcome variable to model as well as reference categories for categorical predictor variables. Further, it runs sequentially by first producing results from simple logistic regression from which the user can select predictor variables to include into the multiple logistic regression, then combine the results of multiple models into a single table. Apart from including only significant predictor variables, based on global/type3 p-values, the user can also choose to include any other variables deemed important by subject matter experts. This flexibility allows for specification of such variables as confounders or effect modifiers even when they are not statistically significant in the simple logistic model. The final output is then processed into a quality publication-ready table and exported into word processing and spreadsheet programs for use in the publication, or if needed, for further hand editing by the authors.

The user must provide input parameters which are specified in Table 1. Unless stated (optional), the other parameters must be provided so that the macro can execute successfully. The ***outevent*** parameter and reference categories for class variables are case sensitive and must be specified in the case they appear in the data dictionary. All other parameters are mainly dataset variables and may be specified in any case. We use lower case for this demonstration. Validation checks enforce these requirements, simplifying debugging errors in macro invocation. The statistician only interacts with sub-macros 1, 4 and 5 by providing input parameters. If a permanent SAS dataset is to be analyzed, the LIBNAME statement can be used to indicate the path or folder where the dataset is located.

**Table 1:** Input parameters for *%svy_unilogit*, *%svy_multilogit* and *%svy_printlogit* macros

| parameter | Description |
| --- | --- |
| **%svy_unilogit** and **%svy_multilogit** macros |  |
| dataset | name of input dataset |
| outcome | name of dependent binary variable of interest e.g., hiv_status |
| outevent | value label of outcome variable (without quotation) to model e.g., Positive, in the case of modeling Hepatitis A risk factors |
| catvars | list of categorical variables (nominal or ordinal) separated by space |
| class | class statement for categorical predictor variables specifying the baseline (reference) category |
| contvars | list of continuous variables separated by space |
| condition | (optional) any conditional statements to create and or fine-tune the final analysis dataset specified using one IF statement |
| strata | (optional) survey stratification variable |
| cluster | (optional) survey clustering variable |
| weight | (optional) survey weighting variable |
| domain | (optional) domain variable for sub-population analysis |
| print | variable for displaying/suppressing the output table on the output window which takes the values (NO=suppress output, YES=show output) |
| **%svysvy_printlogit** macro |  |
| tablename | short name of output table |
| tabletitle | title of output table |
| outcome & outevent | same as defined in **%svy_unilogit** and **%svy_multilogit** macros |
| outdir | directory for saving output files |
| **%runquit** macro |  |
| syserr | SAS in-build macro variable that checks presence of any system errors |

## Results

### Example: Analysis of NHANES dataset

We demonstrate the use of our macro in the analysis of the 2013-2014 National Health and Nutrition Examination Survey (NHANES), a suite of complex surveys designed to assess the health and nutritional status of adults and children in the United States (U.S.). In brief, the main objectives of the survey were to estimate and monitor trends in prevalence of selected diseases, risk behaviors and environmental exposures among targeted populations, to explore emerging public health issues, and to provide baseline health characteristics for other administrative use (28).

NHANES used a four-stage, stratified sampling design, where counties were selected as primary sampling units (PSUs) using probability proportionate to size (PPS) in the first stage. The second stage involved selecting sections of counties that consisted of a block containing a cluster of households with approximately equal sample sizes per PSU. Dwelling units including households were then selected in the third stage with approximately equal selection probabilities. Individuals within a household were selected in the fourth stage. Stratification was done based on selected demographics characteristics of PSUs. Survey weights were then computed using the various sampling probabilities to account for the complex survey design. The data files are freely available to the public on the NHANES website at: https://www.cdc.gov/nchs/nhanes/Index.htm. See (29) for more details regarding the NHANES survey design and contents.

The dataset (`clean_nhanes`) used in this example includes participants’ socio-demographic characteristics including `riagendr` (Gender), `ridageyr` (Age in years at screening), `ridreth1`, (Race/Hispanic origin), `dmqadfc` (Service in a foreign country), `dmdeduc2` (Education level among adults aged 20+ years), and `dmdmartl` (Marital status). The binary outcome variable is `lbxha` (Hepatitis A antibody test result). The aim of the analysis is to investigate factors associated with a positive test for Hepatitis A antibody among participants aged 20+ years who have served active duty in the U.S. Armed Forces (`dmqmiliz`). Appropriate survey weights `wtmec2yr` (sample weights for participants with a medical examination) were applied. The macros were run with user-defined parameters. The user should explicitly specify the reference category for factor variables and for the binary outcome, as shown in Figure 1.

**Figure 1:**
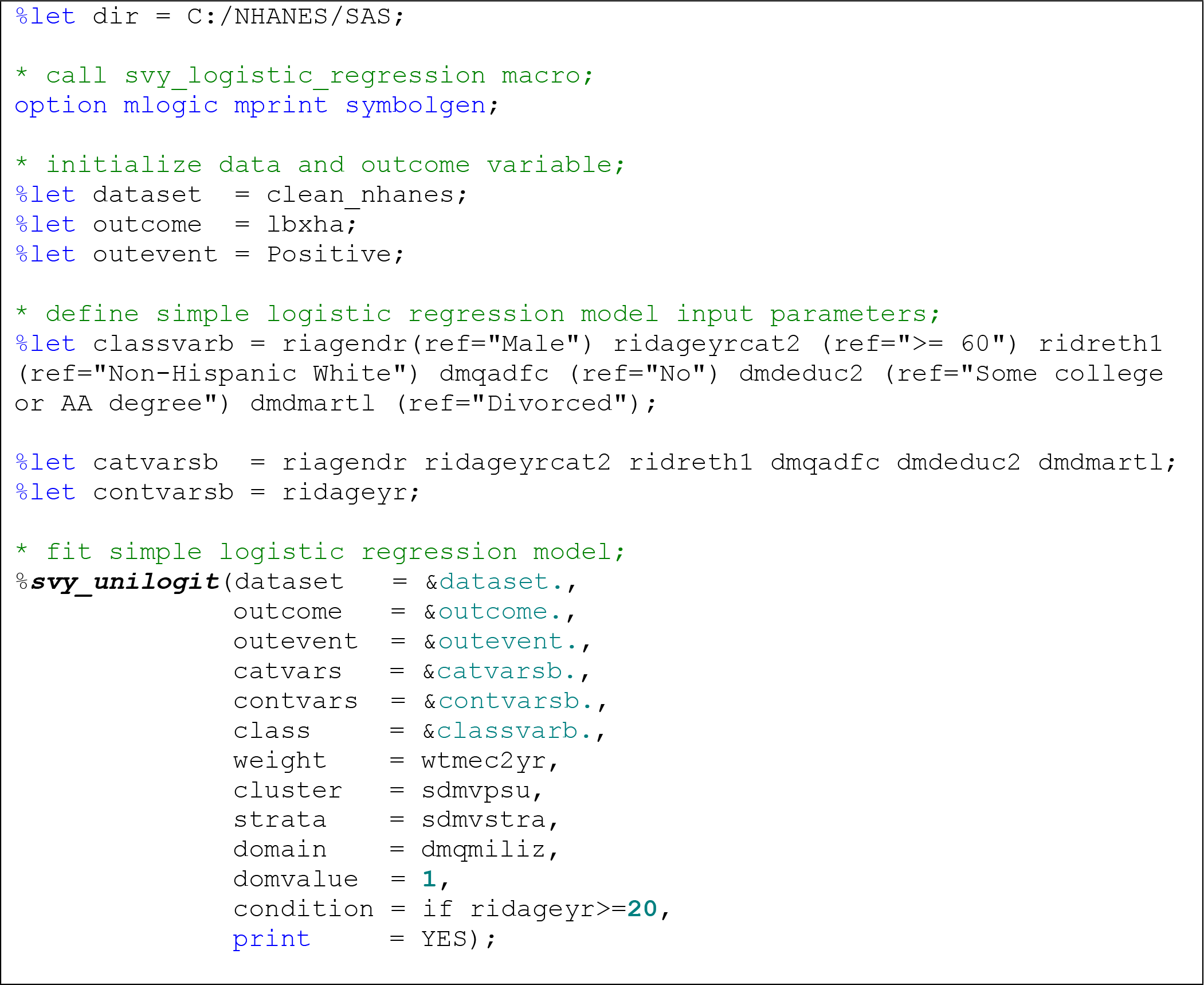
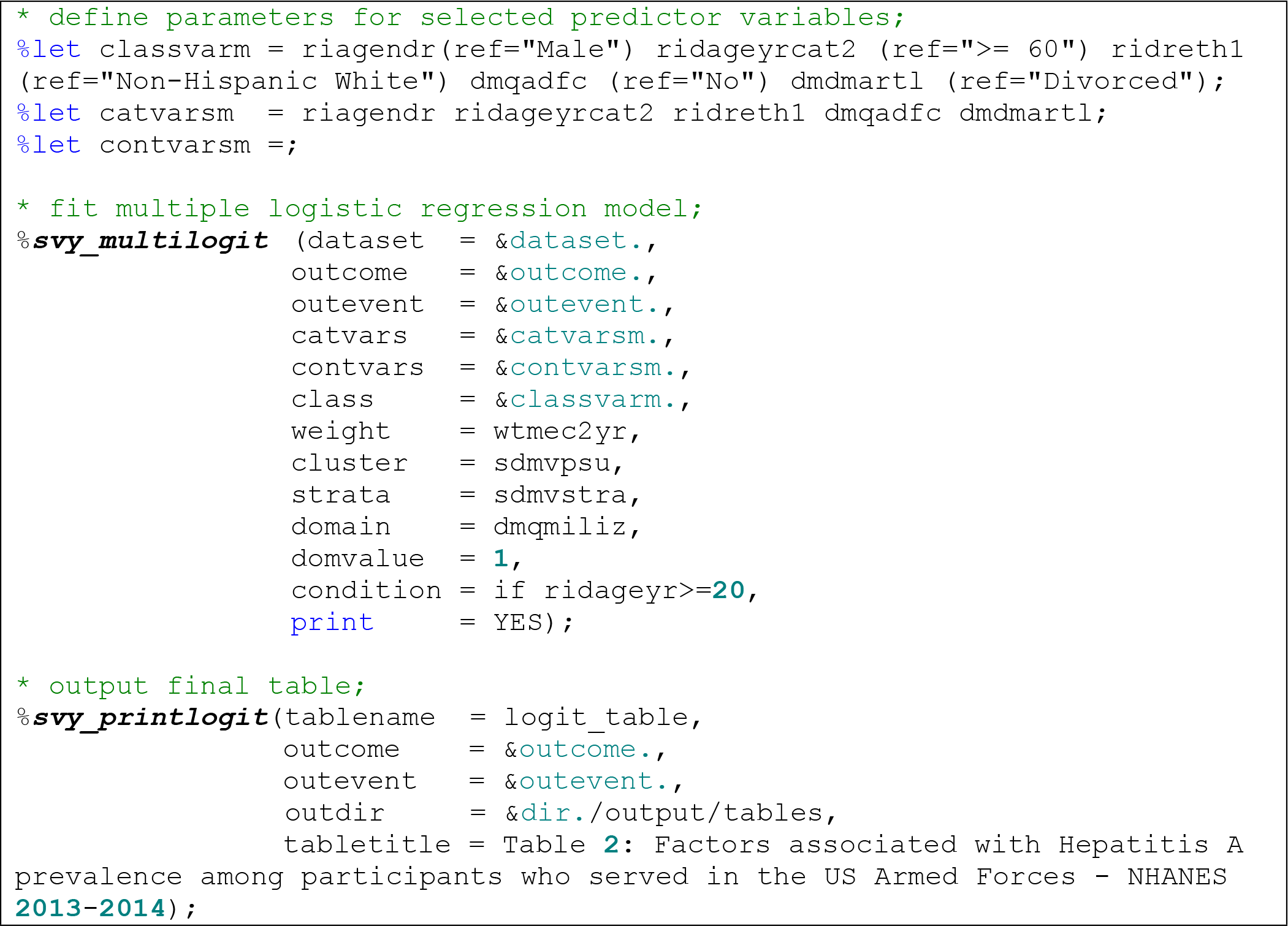
Sample *%svy_logistic_regression* macro call.

The complete SAS output consists of several tables, the majority of which are auxiliary and are used to help in processing the output. Two important output tables are the simple and the multiple logistic regression tables. The simple logistic regression table shows result of bivariate regression as shown in Table 2.

**Table 2:** Output of simple logistic regression model results from *%svy_unilogit* macro

| Factor | N@ | Freq<sup>&</sup> | OR_CI<sup>§</sup> | p_value<sup>α</sup> | g_p_value<sup>β</sup> |
| --- | --- | --- | --- | --- | --- |
| Gender |  |  |  |  |  |
| Male | 473 | 205 (37.7) | ref |  |  |
| Female | 35 | 15 (39.7) | 1.1 (0.4-2.7) | 0.86 | 0.86 |
| Total | 508 | 220 (37.9) |  |  |  |
| Age in years at screening |  |  |  |  |  |
| >= 60 | 322 | 131 (30.7) | ref |  |  |
| 20-39 | 51 | 39 (83.0) | 11.0 (5.4-22.2) | <.01 | <.01 |
| 40-59 | 135 | 50 (31.2) | 1.0 (0.6-1.7) | 0.93 |  |
| Total | 508 | 220 (37.9) |  |  |  |
| Race/Hispanic origin |  |  |  |  |  |
| Non-Hispanic White | 307 | 114 (34.1) | ref |  |  |
| Mexican American | 23 | 14 (67.1) | 3.9 (1.3-12.3) | 0.02 | <.001 |
| Non-Hispanic Black | 126 | 59 (46.1) | 1.7 (1.2-2.4) | 0.01 |  |
| Other Hispanic | 26 | 17 (64.3) | 3.5 (1.0-11.8) | 0.05 |  |
| Other Race | 26 | 16 (52.3) | 2.1 (0.7-6.8) | 0.21 |  |
| Total | 508 | 220 (37.9) |  |  |  |
| Served in a foreign country |  |  |  |  |  |
| No | 243 | 86 (27.4) | ref |  |  |
| Yes | 264 | 134 (48.6) | 2.5 (1.4-4.5) | <.01 | <.01 |
| Total | 507 | 220 (38.1) |  |  |  |
| Education level |  |  |  |  |  |
| Some college or AA degree | 193 | 92 (41.8) | ref |  |  |
| 9-11th grade | 37 | 16 (33.4) | 0.7 (0.4-1.3) | 0.27 | 0.30 |
| College graduate or above | 147 | 51 (32.0) | 0.7 (0.4-1.1) | 0.09 |  |
| High school graduate | 122 | 92 (47.7) | 1.0 (0.6-1.5) | 0.88 |  |
| Less than 9th grade | 9 | 5 (42.6) | 1.0 (0.2-4.5) | 0.97 |  |
| Total | 508 | 220 (37.9) |  |  |  |
| Marital status |  |  |  |  |  |
| Divorced | 75 | 30 (29.4) | ref |  |  |
| Living with partner | 17 | 9 (60.2) | 3.6 (1.1-11.7) | 0.03 | 0.03 |
| Married | 311 | 130 (36.3) | 1.4 (0.8-2.3) | 0.23 |  |
| Never married | 48 | 21 (51.4) | 2.5 (1.1-6.1) | 0.04 |  |
| Separated | 11 | 5 (27.3) | 0.9 (0.3-2.6) | 0.85 |  |
| Widowed | 46 | 25 (44.0) | 1.9 (0.9-4.1) | 0.11 |  |
| Total | 508 | 220 (37.9) |  |  |  |
| Age in years at screening | 508 | 220 (37.9) | 1.0 (1.0-1.0) | <.01 | <.01 |

@ = Total number of observations
& = Frequency of sample cases (and weighted row percentages)
§ = Weighted Odds Ratio (95% confidence interval)
α = Class level p-value
$\beta = \text{Global/Type 3 p-value}$

The table consists of six variables, namely: `Factor` (risk factor variable), `N` (total frequency of observations), `Freq` (frequency of sample cases and corresponding weighted row percentages), `OR_CI` (weighted odds ratio and 95% confidence interval) `p_value` (class level p-value), `g_p_value` (global/type3 p-value). Typically the analyst/researcher selects statistically-important risk factors based on the global/type3 p-values. From this example, all risk factors except gender and education level were statistically significant. However, based on epidemiological considerations, gender and age are often treated as potential confounder variables. Thus they are included in the multiple logistic regression model regardless of statistical significance. Another important aspect to pick from Table 2 is the frequency columns which show the sample size for each factor and each level of the factor. In this example the expected total measurements for each factor was n=508 out of which 220 (37.8%) tested positive for Hepatitis A antibody. All other factors except service in a foreign country (n=507) had complete information available. The importance of this is to ensure that factors with substantive proportion of complete information are selected for inclusion in the multiple logistic regression model. In addition, the row percentages provide guidance on the choice of reference category of factor variables. However, for ordinal factors it is often advisable to use the lowest or highest category as reference, depending on the outcome of interest. After selecting all important variables, the ***%svy_multilogit*** macro is then executed. The ***%svy_printlogit*** macro automatically processes the output into a quality easy to review output as shown in Table 3.

**Table 3:** Quality publication-ready output from the *%svy_printlogit* macro combining results from *%svy_unilogit* and *%svy_multilogit* macros

|  | Hepatitis A<br>antibody <sup>€</sup> |  | Unadjusted |  |  | Adjusted |  |  |
| --- | --- | --- | --- | --- | --- | --- | --- | --- |
| Characteristic | Total<sup>¥</sup> | Positive<sup>£</sup><br>n (%)<sup>&</sup> | OR <sup>ξ</sup> (95% CI)<br><sup>\$</sup> | p-value<sup>α</sup> | Type3<sup>β</sup><br>p-value | OR (95% CI) | p-value | Type3<br>p-value |
| Gender |  |  |  |  |  |  |  |  |
| Male | 473 | 205 (43.3) | ref |  |  |  |  |  |
| Female | 35 | 15 (42.9) | 1.1 (0.4-2.7) | 0.86 | 0.86 | 1.0 (0.3-3.7) | 1.00 | 1.00 |
| Total | 508 | 220 (43.3) |  |  |  |  |  |  |
| Age in years at<br>screening |  |  |  |  |  |  |  |  |
| >= 60 | 322 | 131 (40.7) | ref |  |  |  |  |  |
| 20-39 | 51 | 39 (76.5) | 11.0 (5.4-22.2) | <.01 | <.01 | 13.8 (5.6-34.0) | <.01 | <.01 |
| 40-59 | 135 | 50 (37) | 1.0 (0.6-1.7) | 0.93 |  | 1.3 (0.6-2.8) | 0.54 |  |
| Total | 508 | 220 (43.3) |  |  |  |  |  |  |
| Race/Hispanic origin |  |  |  |  |  |  |  |  |
| Non-Hispanic White | 307 | 114 (37.1) | ref |  |  |  |  |  |
| Mexican American | 23 | 14 (60.9) | 3.9 (1.3-12.3) | 0.02 | <.01 | 3.4 (1.1-10.4) | 0.03 | 0.01 |
| Non-Hispanic Black | 126 | 59 (46.8) | 1.7 (1.2-2.4) | 0.01 |  | 1.9 (1.1-3.2) | 0.02 |  |
| Other Hispanic | 26 | 17 (65.4) | 3.5 (1.0-11.8) | 0.05 |  | 3.4 (0.6-18.6) | 0.16 |  |
| Other Race | 26 | 16 (61.5) | 2.1 (0.7-6.8) | 0.21 |  | 1.4 (0.3-6.9) | 0.65 |  |
| Total | 508 | 220 (43.3) |  |  |  |  |  |  |
| Served in a foreign<br>country |  |  |  |  |  |  |  |  |
| No | 243 | 86 (35.4) | ref |  |  |  |  |  |
| Yes | 264 | 134 (50.8) | 2.5 (1.4-4.5) | <.01 | <.01 | 3.1 (1.9-5.0) | <.01 | <.01 |
| Total | 507 | 220 (43.4) |  |  |  |  |  |  |
| Education level |  |  |  |  |  |  |  |  |
| Some college or AA<br>degree | 193 | 92 (47.7) | ref |  |  |  |  |  |
| 9-11th grade | 37 | 16 (43.2) | 0.7 (0.4-1.3) | 0.27 | 0.30 |  |  |  |
| College graduate or<br>above | 147 | 51 (34.7) | 0.7 (0.4-1.1) | 0.09 |  |  |  |  |
| High school graduate | 122 | 56 (45.9) | 1.0 (0.6-1.5) | 0.88 |  |  |  |  |
| Less than 9th grade | 9 | 5 (55.6) | 1.0 (0.2-4.5) | 0.97 |  |  |  |  |
| Total | 508 | 220 (43.3) |  |  |  |  |  |  |
| Marital status |  |  |  |  |  |  |  |  |
| Divorced | 75 | 30 (40.0) | ref |  |  |  |  |  |
| Living with partner | 17 | 9 (52.9) | 3.6 (1.1-11.7) | 0.03 | 0.03 | 1.8 (0.6-5.1) | 0.27 | 0.02 |
| Married | 311 | 130 (41.8) | 1.4 (0.8-2.3) | 0.23 |  | 1.6 (1.0-2.4) | 0.06 |  |
| Never married | 48 | 21 (43.8) | 2.5 (1.1-6.1) | 0.04 |  | 1.5 (0.9-2.7) | 0.13 |  |
| Separated | 11 | 5 (45.5) | 0.9 (0.3-2.6) | 0.85 |  | 0.9 (0.3-2.9) | 0.88 |  |
| Widowed | 46 | 25 (54.3) | 1.9 (0.9-4.1) | 0.11 |  | 3.2 (1.5-6.7) | <.01 |  |
| Total | 508 | 220 (43.3) |  |  |  |  |  |  |
| Age in years at screening | 508 | 220 (43.3) | 1.0 (1.0-1.0) | <.01 | <.01 |  |  |  |

¥ = Total number of observations
€ = Outcome variable label
£ = Outcome value label of category of interest
& = Frequency of sample cases (and weighted row percentages)
ξ = Weighted Odds Ratio
\$ = Weighted 95% confidence interval for Odds Ratio
α = Class level p-value
β = Global/Type3 p-value

## Discussion

This paper presents an elegant and flexible SAS macro, ***%svy_logistic_regression***, for producing quality publication-ready tables from unadjusted and adjusted logistic regression analyses. Even though a number of SAS macros are available on the internet for processing output from logistic regression into a publication-ready table, they are complex to follow and/or have limited features, thus restricting their adoption. Many macros are not generic and hence can only be used with the data for which they were designed.

The SAS macro presented here is generalized, highly suitable to handle different scenarios, and is simple to implement and invoke from user macros. In addition, our macro includes the row or column total and frequency of prevalent cases of each variable level, which can immediately allow the analyst/researcher to identify levels with sparse data. Row percentages help the researcher in the choice of reference category. Global or type3 p-values shows whether or not a variable is an important predictor. Individual p-values shows if a given variable category is comparable to the reference category. The macro provides validation checks on the input parameters including the dataset, variables and values of variables to ensure that the analyst obtains valid estimates. The output of this SAS macro helps improve efficiency of knowledge generation by reducing the steps required from analysis to clear and concise presentation of results.

## Conclusion

As our contribution to the emerging field of reproducible research, we have provided source code for the SAS macro as well as expected outputs using a publicly available dataset. By publishing this macro, it will allow other SAS macro programmers and users to verify and build upon this code. Production of publication-quality tables is increasingly important as data analyses become more complex, involving larger datasets and requiring more sophisticated computations and tabulation, notwithstanding the need for quick results. This macro helps to make data analysis results readily available, and allows one to publish data summaries in a single document, thus allowing others to easily execute the same code to obtain the same results. The quality, publication-ready results from this macro are suitable for direct inclusion in manuscripts for peer-reviewed journals. The macro can also be used to routinely generate standardized tables. This is especially useful for disease surveillance systems where the same analyses are repeated on a quarterly or annual basis. We hope the published results from this macro will provide

## Author Contribution

JM and SM took part in concept development. JM developed and documented the SAS macro, and prepared the final manuscript. SM tested and debugged the SAS macro. PY helped define user requirements and tested the SAS macro. All authors read and approved of the final manuscript for publication.

## Supporting information

### Supporting results dataset

The NHANES dataset supporting the conclusions of this article is freely available to the public on the NHANES website at: https://www.cdc.gov/nchs/nhanes/Index.htm.

### Supporting software

The source code for this macro is available online at https://github.com/kmuthusi/generic-sas-macros for public access and has been licensed under the terms of the Apache Software License and therefore is licensed under ASL v2 or later. A copy on this license is available at http://www.apache.org/licenses/LICENSE-2.0.html. Sample SAS program call for the macro named *“svy logistic regression anafile.sas”* is provided.

## Acknowledgement

We thank our colleagues in the Surveillance and Epidemiology branch for testing the SAS macro and providing valuable feedback for improvement and also for thoroughly reviewing the manuscript.

